# Cell transformation disrupts the efficiency of chromosome segregation through microtubule detyrosination

**DOI:** 10.1101/246983

**Authors:** Virginia Silió, Jonathan B. Millar, Andrew D. McAinsh

## Abstract

The general principles of chromosome segregation are highly conserved throughout eukaryotic evolution. However, it is unknown whether there are differences in spindle or kinetochore composition or architecture which influence the efficiency chromosome segregation in different cell types. Here we show that the transition of human retinal pigment epithelial cells to a mesenchymal phenotype causes a stabilisation of kinetochore-microtubule attachments and an increase in the frequency of chromosome mis-segregation, due to inefficient error-correction, during mitosis. We find that this is caused by microtubule detyrosination during the epithelial-to-mesenchymal transition and that parthenolide, a tubulin carboxypeptidase inhibitor, efficiently reverts mesenchymal cells to the epithelial mode of chromosome segregation. We propose that reprogramming the post-translational modifications of the mitotic spindle decreases mitotic fidelity and may contribute to CIN in mesenchymal cell populations during tumorigenesis.

---

We recently demonstrated that depletion of the kinetochore null lethal (KNL) 1 protein leads to a dramatic defect in the alignment of chromosomes to the spindle equator (congression) in transformed aneuploid HeLa cells, but only a mild defect in near-diploid immortalized retinal pigment epithelial cells (hTERT-RPE1). Both cell types progress into anaphase with unaligned chromosomes with no delay compared to cells with fully aligned chromosomes, indicative of a defective spindle assembly checkpoint (SAC) ^1^. This is because KNL1 is required for the hierarchical recruitment of Bub1-Bub3, and Mad1-Mad2 checkpoint proteins to kinetochores. These observations raise the intriguing possibility that genetic requirements for chromosome segregation are plastic and may relate to the ploidy and/or transformation state of human cells.

To investigate this, we tested whether chromosome segregation is affected as cells progress through the epithelial to mesenchymal transition (EMT), an event that is important in both normal development, tissue repair and in the initiation of invasive behavior in epithelial cancers ^2^ and generation of circulating tumor cells ^3,4^. Unfortunately, we found that such experiments were not possible with hTERT-RPE1 cells as they already expressed mesenchymal markers, had lost E-cadherin and displayed enhanced migratory behavior, indicating that this cell line is partially transformed (**Fig. S1**). Instead, we turned to the spontaneously arising retinal pigment epithelial 19 cells ^5^ (ARPE19) which have a diploid karyotype, express E-cadherin, lack mesenchymal markers and are non-migratory with the capacity to form monolayers (**Fig. S1**). Treatment of ARP19 cells with transforming growth factor-β1 (TGFβ1) for 6 days caused them to undergo an EMT co-incident with loss of E-cadherin, up-regulation of smooth muscle actin and appearance of migratory behavior, as previously described^6^ (**Fig. S1**).

To monitor chromosome segregation in epithelial and mesenchymal cells we labelled chromosomes with the cell permeable far-red dye (SiR-DNA). We find that the time from nuclear envelope breakdown (NEB) to anaphase onset or completion of chromosome congression is the same in a mesenchymal (TGFβ1 treated) or epithelial cells (normal serum; Fig 1a and b). We did, however, observe an increase in the frequency of lagging chromosomes (from zero in epithelial cells (n=69) to 3.9% in mesenchymal cells (n=51)). This suggests that the process of chromosome bi-orientation in mesenchymal cells is less efficient than in epithelial cells. Following depletion of KNL1, we found that chromosome congression was slower in epithelial cells, with only ~35% of cells completing congression within 60 min (compared to 100% of control cells; Fig. 1b). In contrast, the congression defect was much less severe in mesenchymal cells where ~60% of cells had completed the process within 60 mins (Fig. 1b). While anaphase onset was delayed in epithelial cells, no timing defect was observed in mesenchymal cells compared to *siControl* cells (Fig. 1b). As a result, ~40% of mesenchymal cells entered anaphase with unaligned chromosomes. Importantly, these phenoptypic differences cannot be explained by differences in RNAi efficiency because the levels of KNL1 were reduced to the same extent in both cell types (Fig. 1c).

**Figure 1.**
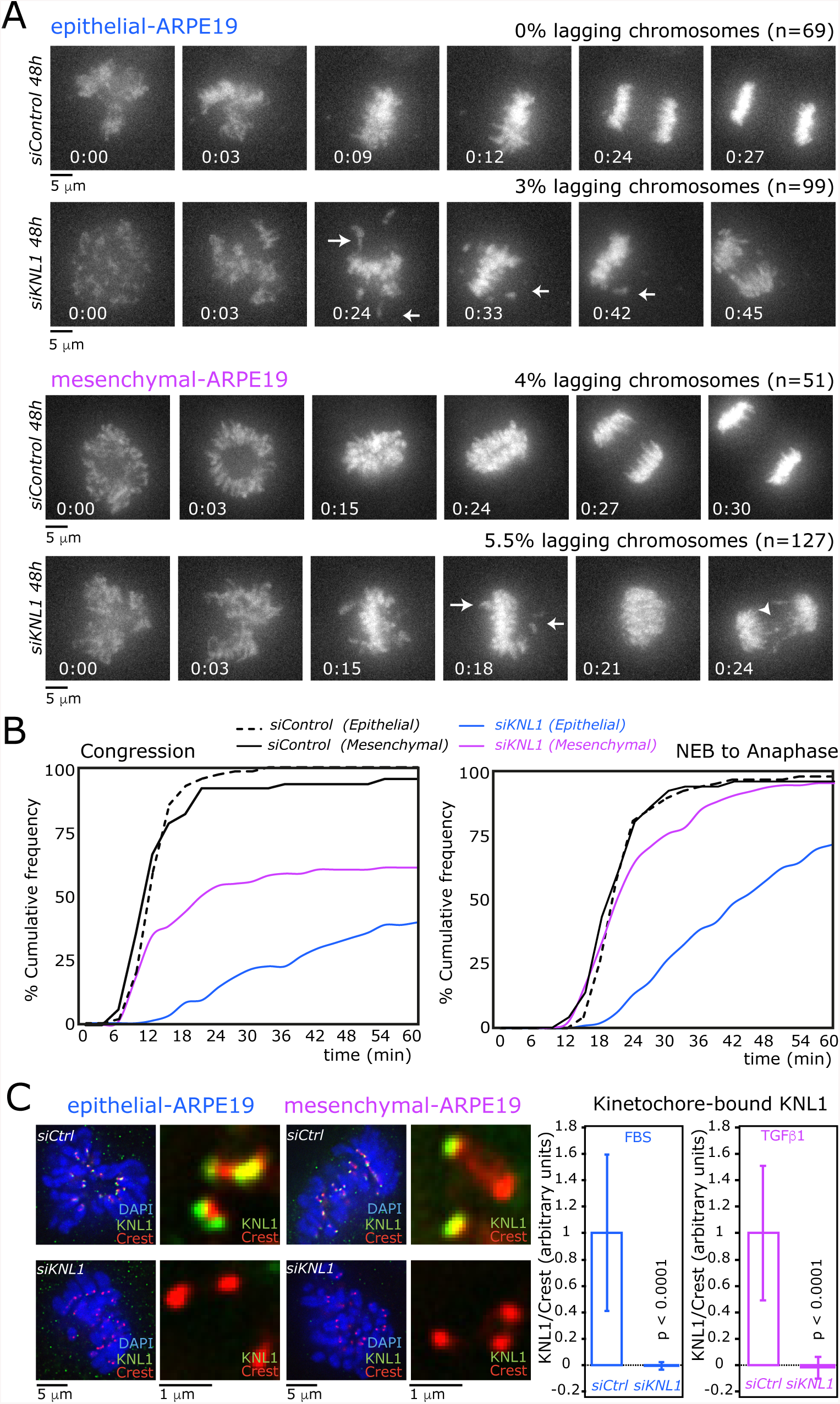
Removal of KNL1 has different phenotype in epithelial and mesenchymal ARPE-19 cells. **(A)** Representative stills of movies of epithelial and mesenchymal ARPE-19 cells treated with the *siCtrl* or *siKNL1*. Arrows show unaligned chromosomes and arrowhead lagging chromosome/chromatid. DNA visualised using SiR-tubulin. Time in min:sec. **(B)** Quantification of the time of congression (NEB to meta-phase) and total time in mitosis (NEB to Anaphase) of cells in A) (epithelial: *siCtrl* N =5, 69 cells, *siKNL1* N= 5, 99 cells; mesenchymal: *siCtrl* N = 5, 51 cells, *siKNL1* N= 5, 127 cells). **(C)** Representative images of immunofluorescence of cells in prometaphase stained for KNL1, Crest and DAPI. Efficiency of the siRNA of KNL1 by quantification of KNL1 relative to Crest intensity at KT, N = 2, 200 KT, data normalized to *siCtrl* media value, p-value from a Student’s *t-*test.

There are two possible explanations for this result. Either KNL1 is only required for spindle assembly checkpoint activation in mesenchymal cells or, secondly, defects in kinetochore microtubule-attachment following loss of KNL1 do not activate the SAC in mesenchymal cells. To investigate this, we treated both epithelial and mesenchymal cells with 330nM nocodazole to generate unattached kinetochores. In either the presence or absence of KNL1, cells were delayed in mitosis suggesting equivalent activation of the spindle assembly checkpoint (**Fig. S2b**). Similar results were found when using lower concentrations of nocodazole that cause only one or two detached kinetochores (**Fig. S2c**). As previously observed, the checkpoint effector Mad2 was recruited to unattached kinetochores, even without detectable Bub1 (as a result of KNL1 depletion; **Fig. S2d**). These data are consistent with our earlier work that showed how unattached kinetochores (in HeLa or RPE1) cells can mount a checkpoint response to unattached, but not unaligned kinetochores with immature microtubule-kinetochore attachment ^1^.

We reasoned that the efficiency of chromosome bi-orientation in epithelial and mesenchymal may be influenced by changes in kinetochore architecture, spindle composition (MAPS or motors) or microtubule dynamics. Although it is well established that the actin cytoskeleton is dramatically remodeled during EMT (e.g. through increased of moesin expression^7^) or the expression of smooth muscle actin (SMA, Figure S1), a mesenchymal marker, much less is known concerning the impact of the EMT on microtubules. Tubulin tyrosine ligase (TTL), which catalyzes the post-translational carboxy-terminal tyrosination of α-tubulin, is down-regulated during EMT^8^. Notably, changes in tubulin tyrosination within the mitotic spindle biases the kinetochore-driven movement of chromosomes towards the spindle equator^9^. To test whether α-tubulin tyrosination is altered during the EMT, we probed extracts from epithelial and mesenchymal cells with antibodies against detyrosinated α-tubulin. We find that treatment with TGFβ causes substantial detyrosination of α-tubulin, consistent with down-regulation of TTL during the EMT (Fig. 2a). We next stained mitotic epithelial and mesenchymal cells with antibodies that recognise either detyrosinated or tyrosinated α-tubulin. This demonstrated that the mitotic spindle is also detyrosinated as cells progress through the EMT (Fig. 2b and **Fig. S2d**). Despite this, other parameters of mitotic spindle function (spindle length, inter-kinetochore distances, kinetochore shape) were unaffected by the EMT (**Fig S3**).

**Figure 2.**
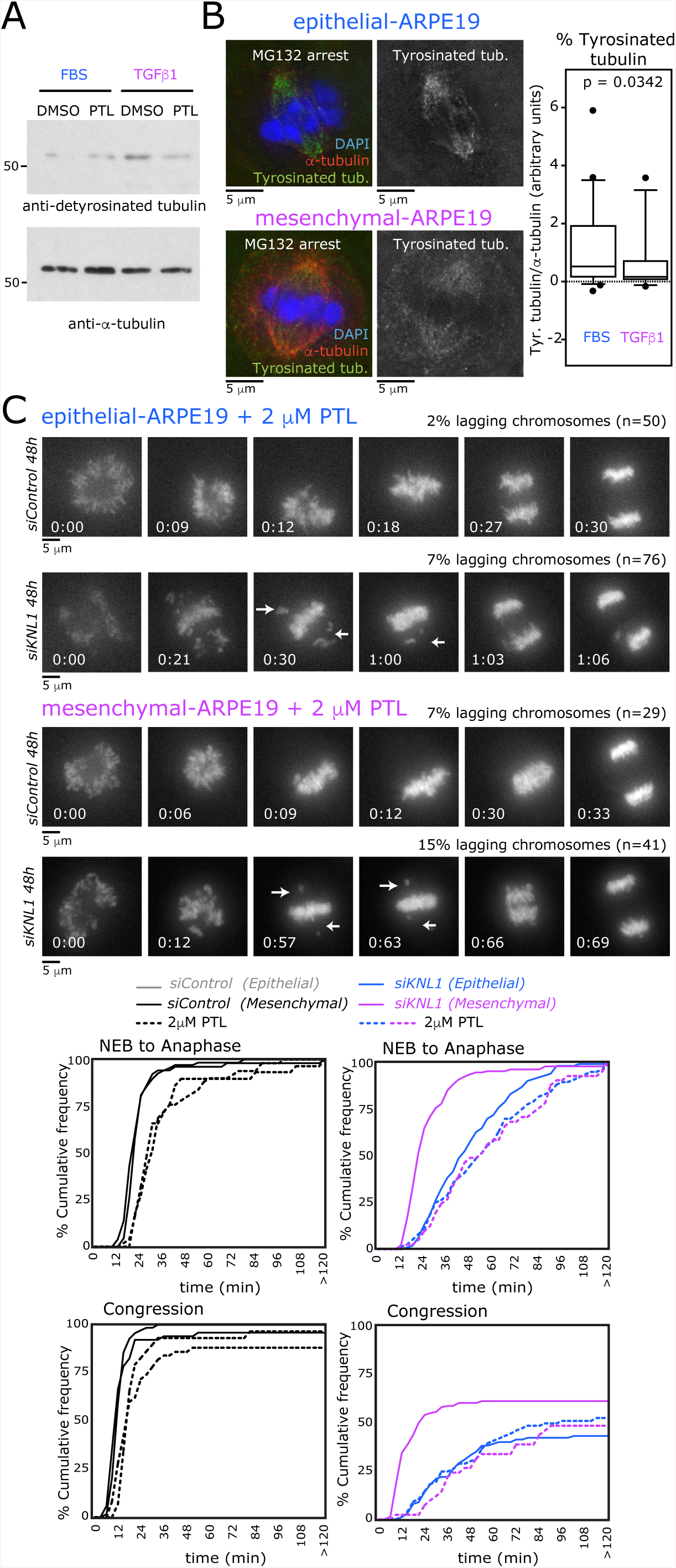
Detyrosination levels can explain the distinct behavior after siKNL1 in epithelial and mesen-chymal ARPE-19 cells. **(A)**Western blot showing the levels of detyrosinated α-tubulin and α-tubulin in epithelial and mesenchymal ARPE-19 cells treated with DMSO (vehicle) or parthenolide (PTL 2 μM).(B) Representative images of ARPE-19 cells arrested in metaphase with MG132 stained for detyros-inated tubulin or tyrosinated tubulin. Quantification of the intensity of the signal at the mitotic spindle of either detyrosinated or tyrosinated tubulin relative to α-tubulin, detyrosinated tubulin, N = 6: epithelial 50 cells, mesen-chymal 37 cells, tyrosinated tubulin, N = 5: epithelial 49 cells, mesenchymal 26 cells, p-value from a Mann-Whitney ney test. **(C)** Representative stills of movies of epithelial and mesenchymal ARPE-19 cells treated with the *siCtrl* or *siKNL1* and parthenolide(PTL 2 μM). Quantification of the time of congression (NEB to metaphase), and total time in mitosis (NEB to anaphase) of these cells. (Epithelial: siCtrl N = 5, 50 cells, *siKNL1* N = 5, 76 cells; mesenchymal: *siCtrl* N = 5, 29 cells, *siKNL1* N= 5, 41 cells). PTL condition is in dotted lines, while results from Figure 1 are shown in straight line for comparison proposes.

Microtubule detyrosination can be partially reversed by treating cells with parthenolide (PTL), which inhibits the tubulin carboxypeptidase responsible for removing tyrosine from α-tubulin, without affecting other post-translational modifications^9–11^ (Fig 2a). In control cells parthenolide had a small effect on the timing of chromosome congression and mitotic timing in both epithelial and mesenchymal cells (Fig. 2c), presumably reflecting the documented role for tyrosination in chromosome congression ^9^. Strikingly, however, although treatment of epithelial cells depleted of KNL1 with parthenolide had no effect, the congression and timing defects in mesenchymal cells were exacerbated, resulting in phenotypes that were indistinguishable from epithelial cells (Fig. 2c). This suggests that the increase detachment rate (and SAC activation) found in epithelial cells can be explained by the synthetic phenotype created by a combination of weakened kinetochores (KNL1 RNAi) and tyrosination of the microtubules. These data imply that kinetochores have a higher affinity for detyrosinated microtubules, which may stabilize inappropriate microtubule-kinetochore (merotelic) attachments. Importantly, PTL treatment did not obviously alter the migratory capacity of mesenchymal cells, which is largely dependent on the actin cytoskeleton, showing that there was no general reversion of the transformation state.

Previous work using TTL knock-out fibroblasts has revealed that tubulin detyrosination inhibits microtubule disassembly and renders microtubule plus-ends resistant to the depolymerase activity of molecular motors^12^. Given that increases in kinetochore-microtubule stability are linked to defects in error-correction^13^ we speculated that transition through the EMT would also lead to mesenchymal cells in which microtubule-attachments are stabilised and error-correction impaired. We first examined the stability of microtubule-kinetochore attachments using a cold-stable assay; stable attachments being more resistant to cold treatment. Indeed, increasing tyrosination of the spindle with parthenolide increases the fraction of mesenchymal cells with destabilized microtubule-kinetochore attachments (Fig. 3a).

**Figure 3.**
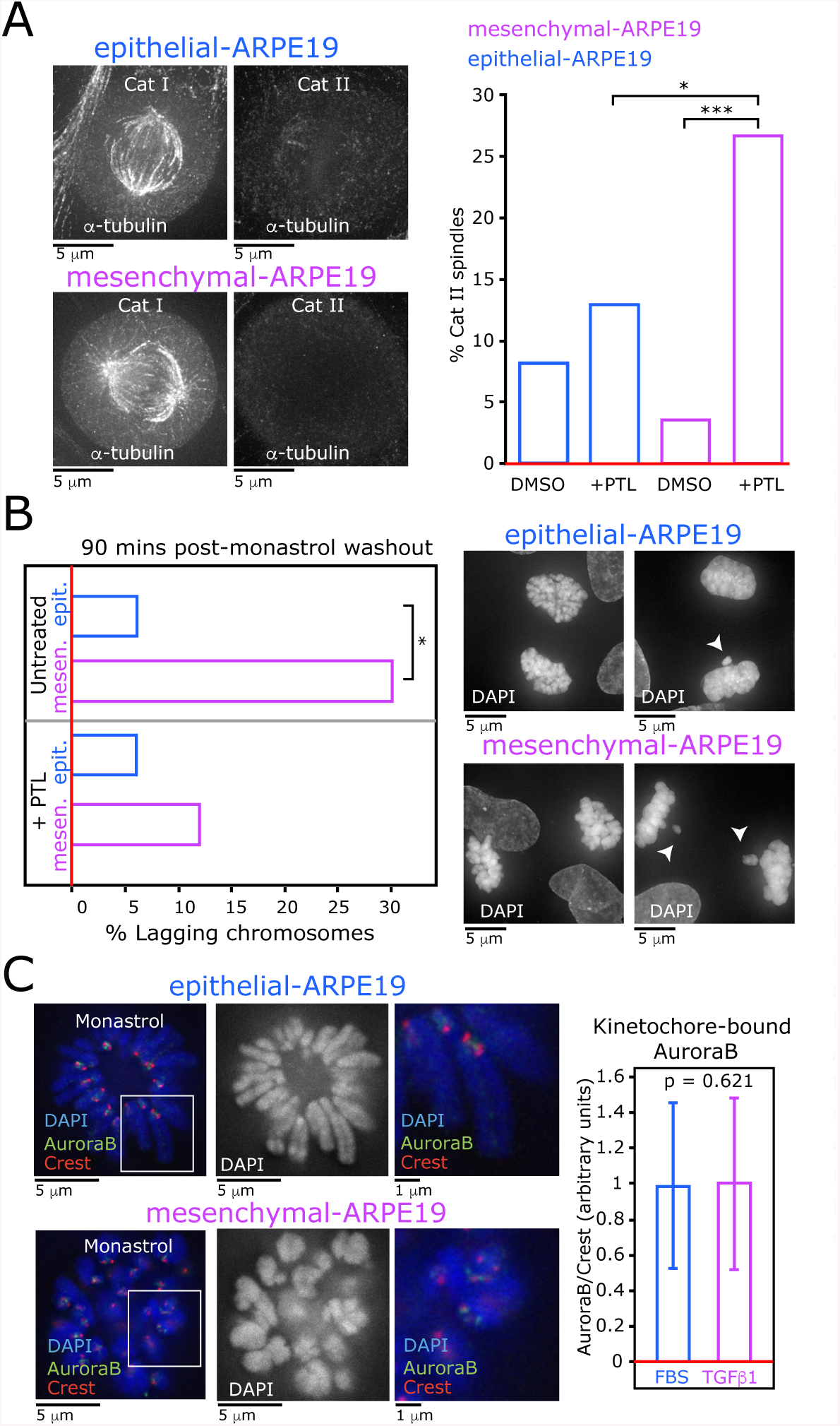
The transition to mesenchymal state increases the probability of chromosome segregation defects. **(A)** Representative images of ARPE-19cells stained for α-tubulin after 5minutes of cold treatment. Number of category I (Cat I) or II (CatII) cells were scored. DMSO N =3, epithelial 61 cells, mesenchymal 57 cells, parthenolide (2 μMPTL) N = 3 epithelial 54 cells,mesenchymal 60 cells. *P <0.05;***P <0.001 (Fisher’s exacttest.**(B)**Percentageofanaphaseandtelophase cells showing alagging chromosome after mon-astrol washout. Untreated N =2, epithelial 83 cells, mesenchymal 88 cells; Parthenolide (2 μMPTL) N = 3 epithelial 258 cells,mesenchymal204cells.***P<0.0001 (Fisher’s exact test **(C)** Representative images of epithelial and mesenchymal ARPE-19 cells arrested with monastrol and stained for Aurora B, Crest and DAPI with quantification of the Aurora B signal intensity relative to Crest in both conditions. N = 3, epithelial 300 kinetochores, mesenchymal 290 kinetochores, data normalized to epithelial ARPE-19 media value, p-value from a Student’s *t*-test.

Next, we tested for defects in error correction by arresting cells with monastrol, a treatment that leads to a monopolar spindle and an increase in syntelic (both kinetochores attached to one spindle pole) and merotelic (one kinetochore attached to both spindle poles) attachments^14^. By releasing the monastrol block, and analysing the cells after 90 minutes, we can assess the efficiency by which cells correct these erroneous attachments and segregate chromosomes without error. In epithelial cells 7.2% of cells made an error in anaphase (lagging chromosome); see Fig. 3b. Notably, there increase in the frequency of errors (to 31.8%) in the frequency of errors in mesenchymal cells. Crucially, the frequency of errors was reduced by ~50% when mesenchymal cells were also treated with parthenolide (Fig. 3b). This defect was not a consequence of altered Aurora-B localization (Fig. 3c), although it was evident that chromosome arms in monopolar spindle were extended in epithelial cells compared to mesenchymal cells. This could be due to more stable microtubules in mesenchymal cells that suppress polar ejector forces (Fig. 3c). Regardless, these data demonstrate that the tyrosination status of microtubules directly impacts the efficiency by which kinetochores can correct erroneous attachments and limit chromosome segregation errors.

In conclusion, our experiments show that the affinity of kinetochores for spindle microtubules is sensitive to the post-translational carboxy-terminal tyrosination of α-tubulin. Tyrosinated microtubules reduce the affinity for the kinetochore, while detyrosination of microtubules increases this affinity. The result is that epithelial cells are more prone to attachment loss (when the kinetochore is perturbed), while error correction in mesenchymal cells is less effective, possibly due to stabilization of inappropriate microtubule-kinetochore attachments, leading to an increased risk of chromosome mis-segregation (Fig. 4). Although the molecular basis for this substrate selectivity is unknown, it is worth noting that the MCAK and Kif2b microtubule depolymerases have a well-established function in error correction and *in vitro* experiments suggest that tubulin detyrosination inhibits MCAK activity ^12,15^.

**Figure 4.**
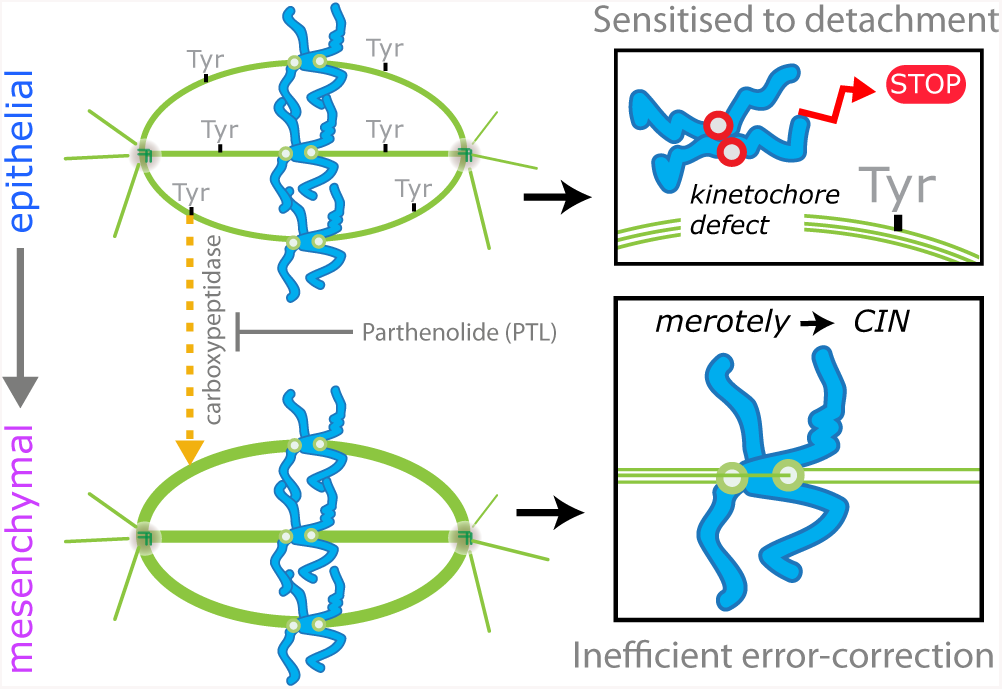
Model for the influence of spindle reprogramming during cellular transformation. The fraction of tyrosinated microtubules within the mitotic spindle is reduced as epithelial cells switch to a mesenchymal phenotype. This leads to increased stability of microtubule-kinetochore attachments. In mesenchymal cells (lower) these stable attachments lead to inefficient error correction and an increased probability of merotelic attachments and chromosome instability. On the other hand, in epithelial cells (upper) the decreased stability of attachments leads to a synthetic phenotype (unattached kinetochores and checkpoint activation) if kinetochores are dysfunctional or mutated.

It is puzzling why the mechanics of chromosome segregation are modulated as cells progress though the EMT. The simplest model is that these changes are simply a by-product of the EMT process, however we cannot rule out that there are advantages to altering the stability of microtubule-kinetochore attachments during development. This is exemplified by the recent finding that the asymmetric tyrosination of meiotic spindles can explain non-Mendelian segregation of maternal and paternal chromosomes ^16^. Tubulin tyrosine ligase (TTL) is frequently suppressed during tumor progression and the resulting tubulin detyrosination is associated with increased tumor aggressiveness, poor prognosis, metastasis and resistance of human cancers to chemotherapeutic drugs ^17–22^. Our experiments suggest that during tumorigenesis the transition to a mesenchymal phenotype and subsequent metastasis are coupled to an increase in chromosomal instability through tubulin detyrosination. Whether modulating the post-translational modifications of tubulin could impact cancer development will be an interesting area of future study.

## Materials & Methods

### Cell culture and drug treatments

ARPE-19 cells were acquired directly from ATCC (CRL-2302) and grown in DMEM/F-12 medium containing 10% fetal bovine serum (FBS), 2.3 gl^-1^ sodium bicarbonate, 100 U/ml penicillin and 100 μg ml^-1^ streptomycin. Only cells at passage 1 and 2 were kept as stocks. All the experiments in this manuscript have been done using passage 2. Cells were maintained at 37°C with 5% CO_2_ in a humidified incubator. Drug treatments were performed at the following concentrations: Nocodazole (SIGMA) 5 nM or 330 nM; monastrol (Tocris) 100 μM; MG132 (12.5 μM, Tocris) 10 μM, Parthenolide (2 μM, SIGMA).

### Epithelial to mesenchymal transition

We designed a protocol from which we started with the same aliquot of cells at the second passage. Cells were first split in two and then and washed twice the following day before incubation with media lacking FBS for 16h to arrest cells in G_0_. Next, cells were switched into media normal media with FBS, or media without FBS but supplemented with 10 ng/ml of TGFβ1 (SIGMA T7039) to induce the epithelial to mesenchymal transition. Experiments were always done with both treatments in parallel following 6 days.

### Depletion experiments

Small interfering (si) RNA oligonucleotides (60 nM); *siControl*^*23*^, *siKNL1*^*24*^, and were transfected with siRNA using oligofectamine (Invitrogen) according to the manufacturer’s instructions. To have the correct amount of cells to transfect, epithelial and mesenchymal cells where re-plated at day 5, keeping the conditioned media until day 6, where they were transfected. After siRNA transfection both conditions were grown in complete media and analyzed 48 h after transfection.

### Live cell experiments

Cells were seeded in a 2-well (Lab-Teck) or 4-well glass bottomed dishes (Greiner Bio One) at day 5 and kept in conditioned media until day 6. Cells were then transfected with siRNA and imaged 48 h before. DNA was followed using SiR-DNA (0.5 μM, Spirochrome) preincubated for 2 h before imaging. We tested this concretration of SiR-DNA in RPE1 cells, and the labeling did not alter the mitotic timing compared to our previous data in RPE1-Histone2B-RFP cells. Movies were acquired on an Olympus DeltaVision Elite microscope (Applied Precision, LLC) equipped with a CoolSNAP HQ2 camera (Roper Scientific) and a stage-top incubator to maintain cells at 37°C and 5% CO_2_. Temperature was stabilized using a microscope enclosure (Weather station; Precision Control) held at 37μC. Image stacks (7× 0.2 μm **optical sections; 1×1 binning) were acquired every 3 minutes for a 12 h period using** a 40x oil-immersion 1.3 NA objective.

### Monastrol release

At day 6, cell were treated with monastrol for 4 h, washed once for fresh media and fixed with PTEM-F after 90 mins. In the PTL condition, cells were kept with PTL during the arrest and the release. Cells were stained with α-tubulin antibodies and DAPI.

### Cold assay

Day 6 cells were treated with MG132 for 4 h, and then incubated on ice for 5 min followed by fixation with cold methanol for 5 min at −20°C. Cells were stained with α-tubulin as described below.

### Immunoblotting

Whole cell lysates were prepared using NP40 10% lysis buffer (50 mM TrisHCl pH 7.6, 150 mM NaCl, 0.5 mM EDTA, 1% NP40) during 40 min on ice. After that, lysates were centrifugated at 14,000 rpm during 30 min and protein concentration in the supernatant was quantified was determined by measurement of absorbance at 280nm using the Protein Assay Dye Reagent Concentrate (Bio-Rad). Lysate was mixed with Laemmli sample buffer (Bio-Rad) and incubated at 98°C for 5 min. Primary antibodies used were rabbit anti-Fibronectin (1:100, SIGMA), anti-E-Cadherin (1:100, Cell Signalling), anti-SMA (1:250, SIGMA), anti-detyrosinated tubulin (1:500, EMD Millipore), and anti-α-tubulin (1:10,000, Sigma), and secondary antibodies anti-mouse and anti-rabbit Horseradish Peroxidase linked raised in donkey (1:10,000, GE Healthcare).

### Immunofluorescence

For immunofluorescence cells were grown on coverslips previously washed with 70 % ethanol and media. Cell were fixed at room temperature (RT) in PTEMF (20 mM PIPES, 10 mM EGTA, 1 mM MgCl_2_, 0.2 % Triton X-100, 4% formaldehyde) for 10 min for KNL1, Mad2, Bub1, Crest, Fibronectin; and fixed with methanol 100% 5 min at −20°C for SMA and tubulin antibodies. Primary antibodies used were mouse anti-Fibronectin (1:100, SIGMA), anti-SMA (1:100, SIGMA), anti-KNL1 (1:500,Abcam), anti-detyrosinated tubulin (1:50, EMD Millipore), anti-tyrosinated tubulin (1:100, EMD Millipore) anti-Bub1 (1:500, Abcam), rabbit anti-Mad2 (1:500, Covance), human anti-Crest (1:200, Antibodies Incorporated), mouse anti-α-tubulin (1:1000, Sigma). Secondary antibodies were goat anti-rabbit/human/mouse/rat conjugated to AlexaFuor-488, 594 or 647 (1:500; Invitrogen). DNA was visualized with DAPI (SIGMA). Three-dimensional image stacks were acquired (1×1 binning) in 0.2 µm steps using a 100× oil NA 1.4 objective on an Olympus Deltavision Elite microscope (Applied Precision, LLC) equipped with a DAPI–fluorescein isothiocyanate– Rhod/TR-CY5 filter set (Chroma) and a Coolsnap HQ2 camera. Deconvolution of image stacks and quantitative measurements were carried out with SoftWorx (Applied Precision, LLC). Kinetochore signals (KNL1, Bub1, Mad2, AuroraB and Crest) were measured in a 10×10 pixel followed by subtraction of background (cytoplasm) and then normalised to the CREST signal (minus background) on the same kinetochore. Modified tubulin signal in the mitotic spindles were quantified in a 19×19 pixel followed by subtraction of background (cytoplasm) and then normalised to the α-tubulin signal in the same place (minus background).

### Statistical analysis

P-values from the figures were obtained using a two-tailed unpaired test with 95% of confidence, except for figure 2b were we used a two-tailed Mann-Whitney test, and in figure 3a and b where we used Fisher’s Exact test. Number of cells and experiments are detailed in each figure legend.

## Acknowledgements

This work was supported by MRC program grant MR/K001000/1 to A.D.M. and J.B.M. A.D.M is also supported by a Wellcome Trust Senior Investigator Award (grant 106151/Z/14/Z) and a Royal Society Wolfson Research Merit Award (grant WM150020).

## Author contributions

All experiments and data analysis was carried out my V.S. Interpretation of data and project planning was carried out by A.D.M, V.S. and J.B.M. A.D.M wrote the manuscript with input from V.S. and J.B.M.

## Author information

The authors declare no competing financial interests.

